# Pf-HaploAtlas: an interactive web app for spatiotemporal analysis of *P. falciparum* genes

**DOI:** 10.1101/2024.07.16.603783

**Authors:** Chiyun Lee, Eyyüb S. Ünlü, Nina F.D. White, Jacob Almagro-Garcia, Cristina Ariani, Richard D. Pearson

**Affiliations:** Wellcome Sanger Institute, Hinxton, UK

**Author notes:** Correspondence should be addressed to R.D.P. These authors contributed equally to this work (Chiyun Lee, Eyyüb S. Ünlü, Nina F.D. White).

## Abstract

Monitoring the genomic evolution of *Plasmodium falciparum* - the most widespread and deadliest of the human-infecting malaria species - is critical for making decisions in response to changes in drug resistance, diagnostic test failures, and vaccine effectiveness. The MalariaGEN data resources are the world’s largest whole genome sequencing databases for *Plasmodium* parasites. The size and complexity of such data is a barrier to many potential end users in both public health and academic research. A user-friendly method for accessing and interpreting data on the genetic variation of *P. falciparum* would greatly enable efforts in studying and controlling malaria. We developed Pf-HaploAtlas, a web application enabling exploration of genomic variation without requiring advanced technical expertise. The app provides analysis-ready data catalogues and visualisations of amino acid haplotypes for all 5,102 core *P. falciparum* genes. Pf-HaploAtlas facilitates comprehensive spatial and temporal analyses of genes and variants of interest by using data from 16,203 samples, from 33 countries, and spread between the years 1984 and 2018. The scope of Pf-HaploAtlas will expand with each new MalariaGEN *Plasmodium* data release. Pf-HaploAtlas is available online for public use at https://apps.malariagen.net/pf-haploatlas, which allows users to download the underlying amino acid haplotype data, and its source code is freely available on GitHub under the MIT licence at https://github.com/malariagen/pf-haploatlas.

## Introduction

Malaria presents a significant global health challenge. In 2023, the World Health Organization (WHO) reported an estimated 249 million malaria cases across 85 countries, and 608,000 fatalities from the disease (WHO 2023). Over three-quarters of these deaths occurred in children under five years old. Malaria-causing parasites can exhibit rapid genomic adaptation to control measures (WHO 2023). For example, genetic mutations have arisen in certain parasite genes, resulting in drug resistance, which makes malaria harder to treat. Genomic surveillance can identify these changes and inform on how best to respond. It has been successfully applied to monitor the evolution of the deadliest malaria parasite, *Plasmodium falciparum* (MalariaGEN et al., 2023). Genomic surveillance by the GenRe-Mekong project across eight countries tracked drug-resistant *P. falciparum* parasites, which helped spot their spread into new areas like Vietnam and Laos, guiding decisions for National Malaria Control Programs (Jacob et al., 2021). It is crucial to monitor such known genetic mutations, but also critical to search more widely for as-yet unstudied drug resistance mutations, which may gain importance over time. Other evolutionary threats to malaria control and elimination are also important to consider. Since 2021 two new malaria vaccines have been approved - RTS,S and R21 - and these may increase pressure for *P. falciparum* to make evolutionary adaptations (WHO 2023). All these changes prompt an increased urgency for robust, timely, ubiquitous genomic surveillance of *P. falciparum*.

Achieving such comprehensive surveillance, however, faces several challenges. Processing and interpreting genomic sequence data requires specialist computing infrastructure and technical expertise, which are not universally available (Hill et al., 2023). There is a need for tools which address these barriers and increase the collective power of the community to perform genomic surveillance. To this end, we have developed a user-friendly web app, Pf-HaploAtlas, that affords anyone with an internet connection the ability to study genetic mutations across any gene in the *P. falciparum* core genome.

## Methods

Pf-HaploAtlas currently uses data generated using the publicly available MalariaGEN Pf7 whole genome sequencing data release (MalariaGEN et al., 2023) and will be updated once new data sets are made publicly available. Only *P. falciparum* samples which have passed quality control were included in the generation of Pf-HaploAtlas outputs, totalling 16,203 samples. To ensure Pf-HaploAtlas runs quickly, graphical outputs are generated from precomputed data files which contain haplotype information for all samples. Haplotypes were computed using custom Python scripts to apply genotypes from the GT field of the Pf7 VCFs to the 3D7 reference sequence, and then translate the nucleotide sequence into a) amino acid sequence and b) lists of non-synonymous variants. We used all SNPs that passed filtering. For multi-exon genes we concatenated nucleotide sequences from each exon before translating to amino acid haplotype. We excluded any haplotype where any genotype call was missing, or where the resulting amino acid haplotype was heterozygous or included a premature stop codon, so only clonal gene haplotypes were included in Pf-HaploAtlas. We calculated haplotypes for each gene within the core genome, which excludes multigene families in the sub-telomeric and internal VAR regions (Miles et al. 2016). Only the first isoform of each gene according to PlasmoDB was used in analysis for genes with multiple isoforms. Comprehensive data catalogues of sample- and population-level haplotype data can be downloaded from the app in the “Download data” section.

Pf-HaploAtlas was built using Streamlit - an open-source Python-based frontend framework. At the time of writing, the app is deployed using the Streamlit community cloud, though in future the method of deployment may change and hence, we recommend using the stable link to the app: https://apps.malariagen.net/pf-haploatlas.

## Results

Pf-HaploAtlas provides downloadable data catalogues and visualisations of haplotypes for all 5,102 core genes by using data from 16,203 samples, from 33 countries, and spread between the years 1984 and 2018, facilitating comprehensive spatial and temporal analyses of genes and variants of interest.

A typical user workflow is as follows:

1. The user can search for, or scroll through the list of core genes in a dropdown menu. Genes are identified by their unique gene ID and, where applicable, their gene name as used on PlasmoDB (Alvarez-Jarreta et al. 2023). Upon selection of a gene, the following plot for the selected gene is displayed:
2. The Haplotype UpSet plot consists of three panels which provide an overview of the haplotypes present for each gene (Figure 1A). For each unique haplotype, the UpSet plot shows the total number of samples (top panel), the proportion of populations making up the haplotype (middle panel), and the mutation make-up of the haplotype (bottom panel). Unique haplotypes are displayed in order of decreasing prevalence from left to right. The populations shown in the middle panel bar plot correspond to the ten major subpopulations defined in Pf7 (MalariaGEN et al., 2023). These are colour-coded for easy interpretation, and actual values may be viewed upon hovering the cursor over each bar. On the x-axis of these barplots, *P. falciparum* lab strains (e.g., 3D7, 7G8, Dd2) with matching haplotype are shown. Investigation into a specific haplotype can be done by clicking on any of the data elements of the haplotype, e.g., clicking on the bar charts or mutations of the haplotype. Upon selection of a haplotype, the following two plots for the selected haplotype are displayed:
3. The abacus plot displays haplotype frequencies (proportion of samples with the selected haplotype) across locations and time (in years) (Figure 1B). The colour intensity of each “bead” on the abacus plot corresponds to the haplotype frequency in each year and in each location (at an administrative division level). Haplotypes at fixation are marked with “100” to highlight 100% frequency, whilst “beads” with 0% haplotype frequency are crossed out.
4. The world map plot displays the haplotype frequency over a user-defined interval of time (in years) at a country-level on a global map (Figure 1C).

Finally, the user is able to access more information, modify the data filtering, and download the raw data by clicking the “Click to see more about the data” tab.

**Figure 3:**
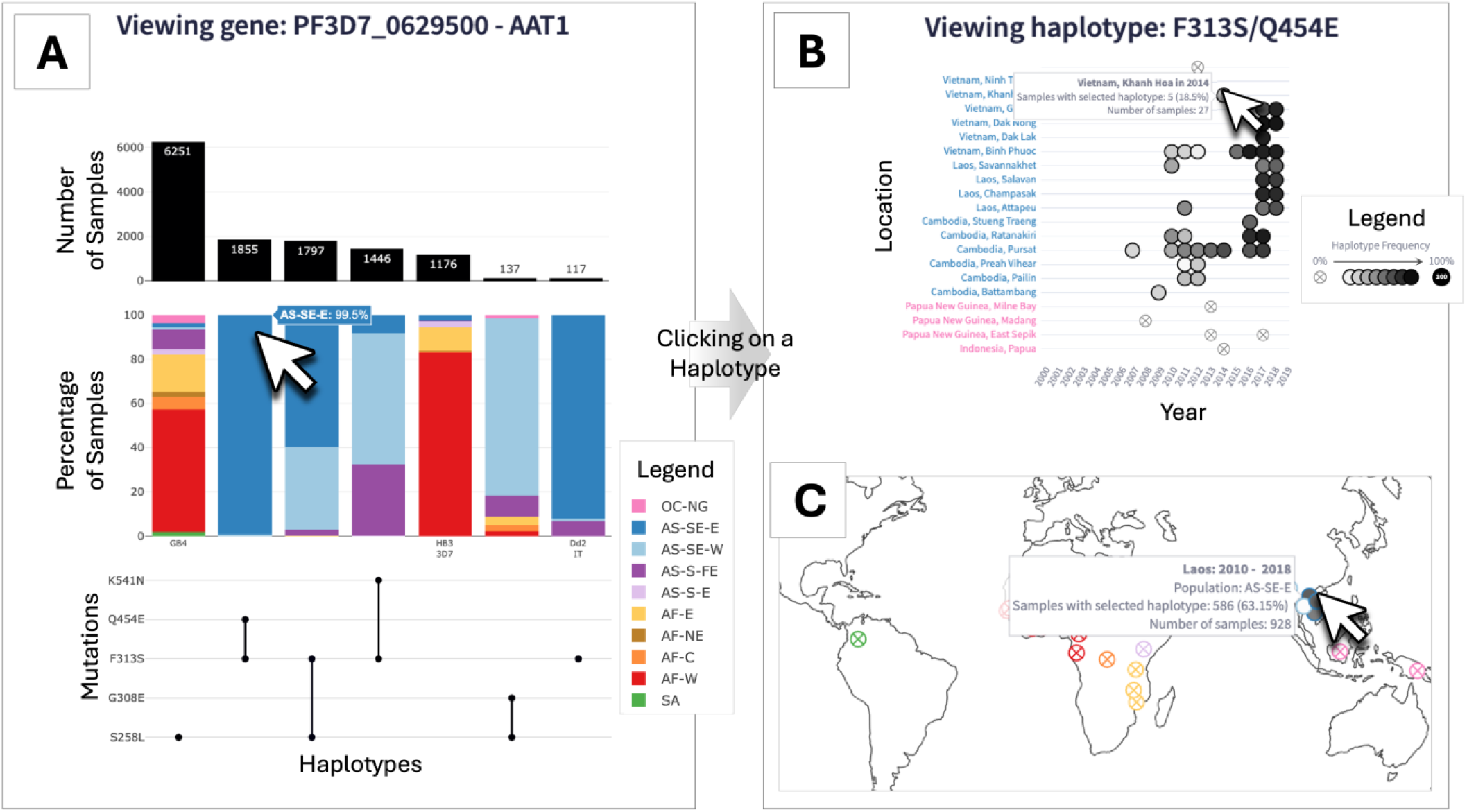
Overview of the Pf-HaploAtlas app, and demonstration of its use for investigating *aat1*, a gene that has recently become of interest in malaria genomic surveillance. **(A)** Haplotype UpSet plot for *aat1*. The left-most haplotype, S258L, is the most prevalent and shows wide geographic distribution, as depicted by the wide variety of subpopulation colours. The cursor hovers over the second most frequent haplotype, F313S/Q454E, which is almost exclusively found in the eastern part of southeast Asia (AS-SE-E). Clicking on this haplotype generates the abacus plot **(B)** and the world map plot **(C)** directly below the Haplotype UpSet plot within the app. The abacus plot for F313S/Q454E highlights an evident increase in frequency in several locations across Laos, Cambodia, and Vietnam over the past two decades, as depicted by the gradient of increasing colour intensity from left to right **(B)**. All images of plots have been edited to retain visual clarity within the manuscript. With all plots, hovering over data elements with the cursor provides more information.

### Use-case demonstration: following trends in compensatory mutations in *aat1* and drug resistance haplotypes of *crt*

A key use case of Pf-HaploAtlas is the discovery of emerging trends in relatively understudied genes. For example, selecting the putative amino acid transporter gene, *aat1*, gives insights into the growing frequency of certain haplotypes linked to drug resistance within the Greater Mekong Subregion over recent years. *aat1* has only very recently been associated with resistance to chloroquine, as a compensatory mutation for the drop in fitness caused by drug-resistant chloroquine resistance transporter gene (*crt*) haplotypes (Amambua-Ngwa et al., 2023), and is therefore unlikely to be included in targeted amplicon panels designed before 2023 for genomic surveillance. However, the whole genome data used by Pf-HaploAtlas enables this gene to be studied. From the Haplotype UpSet plot, it is evident that the most common *aat1* haplotype has a wide geographic distribution, but the second most frequent haplotype, F313S/Q454E, is almost exclusively found in the eastern part of southeast Asia (Figure 1A). The abacus plot for F313S/Q454E highlights an evident increase in frequency in several locations across Laos, Cambodia, and Vietnam over the past two decades, as depicted by the gradient of increasing colour intensity from left to right. This is an example of the app being used to identify possible signs of positive selection (Figure 1B).

Alongside lesser-known genes, Pf-HaploAtlas is also able to highlight trends in haplotypes of key drug-resistance genes. The *crt* gene has been associated with chloroquine resistance for over two decades and remains an important locus for tracking evolutionary threats to malaria control measures (Djimdé et al. 2001; Hagenah et al. 2023). The breadth of data included in Pf-HaploAtlas allows users to discover important recent genetic changes. For example, users can identify the accumulation of recently emerged *crt* mutations on top of the widespread Dd2 resistance haplotype, such as T93S, H97Y, F145I, and I218F (Hamilton et al. 2019). These haplotypes are associated with resistance to the antimalarial drug piperaquine and can be seen emerging in Vietnam, Cambodia, and Laos after 2010 using the Pf-HaploAtlas (Supplementary Figure 1).

## Concluding remarks

Pf-HaploAtlas is a significant addition to the field of malaria genomics. By leveraging the largest openly available whole genome dataset for *P. falciparum*, Pf-HaploAtlas serves as a database of analysis-ready haplotype data, as well as providing detailed interactive visualisations of spatiotemporal haplotype diversity for thousands of samples.

Pf-HaploAtlas has several advantages over existing tools for malaria genomic surveillance. The use of whole genome data enables Pf-HaploAtlas users to explore all 5,102 core genes of *P. falciparum*. This vastly increases the potential for making new discoveries of evolutionary trends in previously understudied genes. Even within well-known genes associated with malaria control measures, Pf-HaploAtlas represents a step forwards thanks to its range of spatiotemporal visualisations and aggregation functions. The Haplotype UpSet plot is unique amongst currently available malaria surveillance tools in its visualisation of haplotype frequencies across global populations. Pf-HaploAtlas’ abacus plot enables tracking of haplotype frequencies across time and multiple locations – a crucial factor for comparative analysis of evolution between areas with different malaria control programmes. These features collectively provide the most granular and comprehensive public tool for *P. falciparum* genomic surveillance, facilitating the identification of important trends and guiding timely, effective interventions in malaria control efforts.

Pf-HaploAtlas’ methodology is not limited to *P. falciparum* and could readily be adapted to other organisms requiring genomic surveillance. To our knowledge, no existing tool showcases haplotype frequencies. As shown in the Pf-HaploAtlas, spatio-temporal visualisation for haplotypes is straightforward to interpret, making it accessible to a broader audience, including those who may not be specialists in evolutionary genetics.

Aside from its analytical approach and data visualisation, Pf-HaploAtlas is an important contribution to open-source genomics specifically. The app’s foundation on open data and open-source software makes it accessible to a broad range of users interested in genomic surveillance. Building tools with Streamlit (https://streamlit.io/), which has a relatively shallow learning curve compared to standard web development frameworks, greatly lowers the barrier to entry for data scientists and bioinformaticians who wish to publish their surveillance tools for public use in the form of an app. Combined with providing an analysis-ready database of genetic variation freely available for download, Pf-HaploAtlas sets an example for democratising and decentralising genomic data analysis, empowering users worldwide to contribute to surveillance efforts.

To maintain its utility and expand its scope, future iterations of Pf-HaploAtlas will be updated with the newest releases of *P. falciparum* whole genome data. The proportion of missing data across space and time would be reduced if there is a move towards more systematic, longitudinal sampling and whole genome sequencing of *Plasmodium* spp. in the malaria genomic surveillance community. Subsequent versions of Pf-HaploAtlas could be expanded to cover other malaria parasites, such as P. vivax, which contributed to 6.9 million malaria cases and 3% of the global malaria burden in 2022 (WHO 2023). With continuous improvements to data integration and releases of new functionalities, Pf-HaploAtlas has the potential to remain at the forefront of malaria genomic surveillance tools.

## Acknowledgements

Pf-HaploAtlas currently uses data generated using the MalariaGEN Pf7 data release which was made possible by clinical parasite samples contributed by partner studies, whose investigators are represented in the data release’s author list.

## Author contributions

R.D.P. conceptualised the idea for this work. C.L. carried out the project administration and designed the technical architecture. C.L., E.S.Ü., R.D.P. and N.F.D.W. developed the software. C.L., E.S.Ü., and N.F.D.W. drafted the manuscript. E.S.Ü. and N.F.D.W. performed data curation. J.A.G., C.A., and R.D.P. supervised the project and reviewed the manuscript.

## Data availability

Pf-HaploAtlas is available online for public use at https://apps.malariagen.net/pf-haploatlas, and its source code is freely available on GitHub under the MIT licence at https://github.com/malariagen/pf-haploatlas.

## Supplementary material

**Figure S1:**
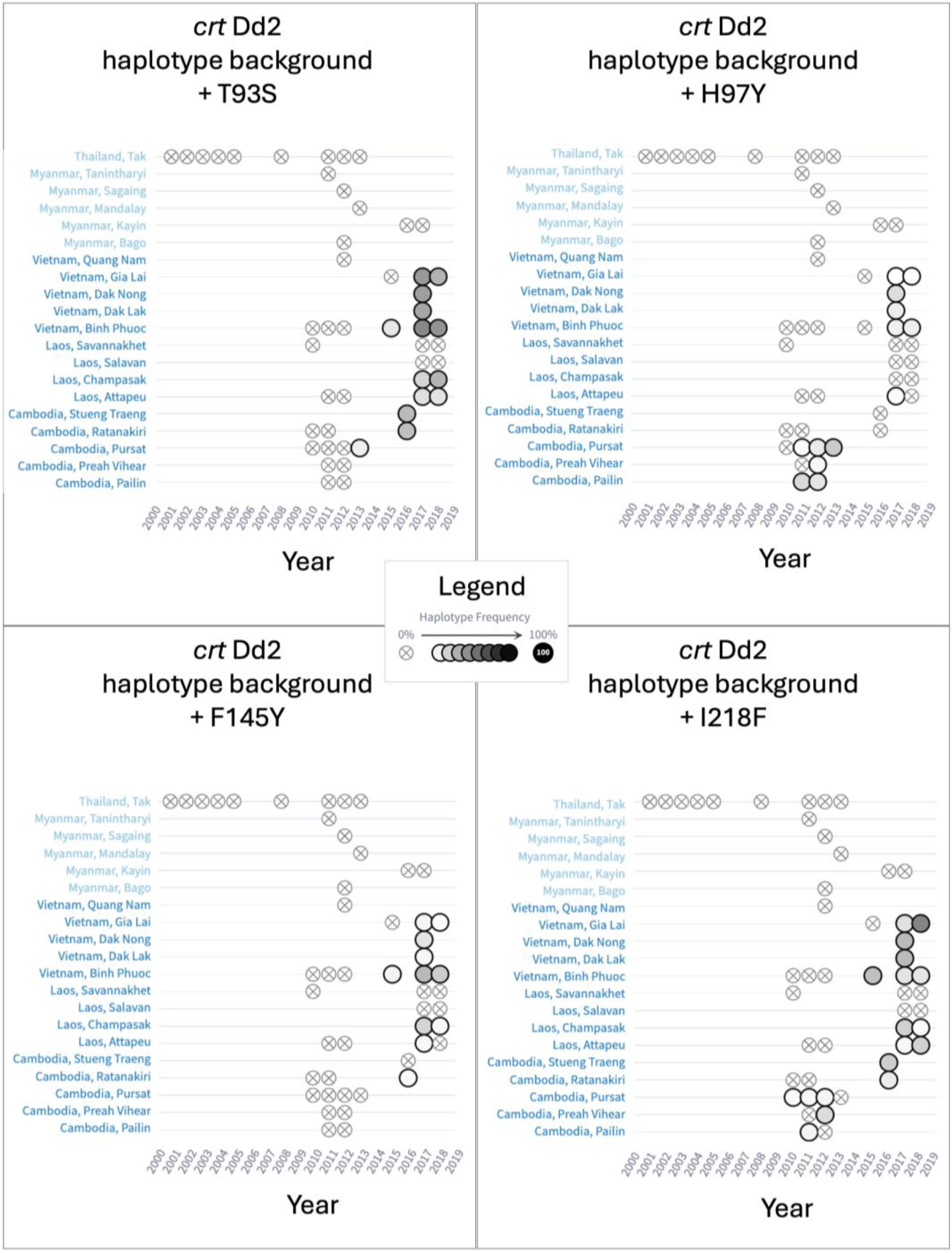
Abacus plots of various haplotypes of interest in the *crt* gene.

## References

1. Alvarez-Jarreta J, Amos B, Aurrecoechea C et al. VEuPathDB: the eukaryotic pathogen, vector and host bioinformatics resource center in 2023. Nucleic Acids Research 2024;52:D808–16.

2. Amambua-Ngwa A, Button-Simons KA, Li X et al. Chloroquine resistance evolution in Plasmodium falciparum is mediated by the putative amino acid transporter AAT1. Nat Microbiol 2023;8:1213–26.

3. Djimdé A, Doumbo OK, Cortese JF et al. A molecular marker for chloroquine-resistant falciparum malaria. N Engl J Med 2001;344:257–63.

4. Hagenah LM, Dhingra SK, Small-Saunders JL et al. Additional PfCRT mutations driven by selective pressure for improved fitness can result in the loss of piperaquine resistance and altered Plasmodium falciparum physiology. mBio 2024;15:e01832–23.

5. Hamilton WL, Amato R, Van Der Pluijm RW et al. Evolution and expansion of multidrug-resistant malaria in southeast Asia: a genomic epidemiology study. The Lancet Infectious Diseases 2019;19:943–51.

6. Hill V, Githinji G, Vogels CBF et al. Toward a global virus genomic surveillance network. Cell Host & Microbe 2023;31:861–73.

7. Jacob CG, Thuy-Nhien N, Mayxay M et al. Genetic surveillance in the Greater Mekong subregion and South Asia to support malaria control and elimination. eLife 2021;10:e62997.

8. MalariaGEN, Abdel Hamid MM, Abdelraheem MH et al. Pf7: an open dataset of Plasmodium falciparum genome variation in 20,000 worldwide samples. Wellcome Open Res 2023;8:22.

9. Miles A, Iqbal Z, Vauterin P et al. Indels, structural variation, and recombination drive genomic diversity in Plasmodium falciparum. Genome Res 2016;26:1288–99.

10. World Health Organization. World malaria report 2023. 30 November 2023 (https://www.who.int/teams/global-malaria-programme/reports/world-malaria-report-2023).

